# An Interactive Bidirectional De Novo Protein Design System

**DOI:** 10.64898/2026.10.27.762208

**Authors:** Chance Gammill, Lakshman Tamil

## Abstract

We present a lightweight, interactive server that combines two generative models, a structure-aware graph neural network (GNN) for de novo sequence design and an ESMFold-based structure predictor, into a single bidirectional design loop. The tool requires no software installation beyond a web browser, and it visualizes and quantifies both directions of the sequence-structure mapping between predicted sequences and structures. Through the same interface, an included user-accessible Kabsch alignment tool superimposes a predicted structure onto the target backbone. It reports the resulting root mean square deviation directly in the interface. Rather than a new model, this paper contributes the tool itself: an accessible, flexible, and openly sourced codebase that lowers the barrier to using and analyzing a model. This interactive server can also be used to validate design hypotheses interactively and reused by other researchers with their own checkpoints. While slightly more challenging, the codebase includes a section for users to add their own model structures while still using the interactive server code. This paper describes the system’s architecture, interface, and intended workflow, and discusses its role as an open-source tool. All data and models used in this paper are derived from our previous companion papers [3, 4].

## 1. Introduction and Motivation

In our two previous companion papers, *De novo Protein Generating Model via Structure-Aware Graph Neural Networks* [3] and *Iterative Refinement for Structural Optimization of De Novo Protein Sequences* [4], we evaluated both contributions against held-out structures using batch scripts and aggregate statistics. While that evaluation method is appropriate for establishing model and procedure performance, it is not well designed for single-protein predictions and evaluations. A collaborator without a machine learning background, or a student, would likely struggle to use these metric values meaningfully. This stems from comparing statistics and biology. While a root-mean-square deviation (RMSD) score may make a protein seem structurally similar, a kink in the wrong location or a positive amino acid in place of a negative would make the resulting structure useless, so we designed a tool to help close that gap. Allowing users, no matter the background, to see how closely, at which locations, and with which models their synthesized proteins performed best.

The resulting interface is shown in Figure 1. Overall, this lets a researcher fold a sequence, design a sequence for a hand-built or imported backbone, test multiple solutions, and compare them against the original sequence’s predicted structure without writing code, managing a Python environment, or leaving the browser. Additionally, because structures render as they are generated, users can easily test and visualize hypotheses about how a backbone change, a longer helix, a modified loop, or an imported PDB chain with one region edited by hand affects the designed sequence. For further analysis, a user-accessible Kabsch [6] superposition (Section 4.4) aligns a predicted structure to its target backbone. It reports the resulting C*α* RMSD so that any design can be analyzed both visually and numerically without separate analysis. This tool is released as open-source code; any checkpoint trained with the same backbone-conditioned sequence design architecture, including the checkpoint described in our companion paper [3], but also checkpoints trained by other groups on the same architecture, can be utilized by changing a single path; however, changes to the model architecture will require changing the model config or substituting the model import code. This makes the tool a general-purpose front-end codebase for structure-conditioned sequence design models. Whereas other existing tools are limited to folding or inverse folding: folding tools (e.g. ESM-Fold [8], AlphaFold [5] web interfaces) predict structure from sequence, while inverse folding tools (e.g. the ProteinMPNN [2] web server) predict sequence from structure. Few freely available tools support both folding methods within the same session against the same visual model, which is an important step in a hand-designed protein workflow.

**Figure 1.**
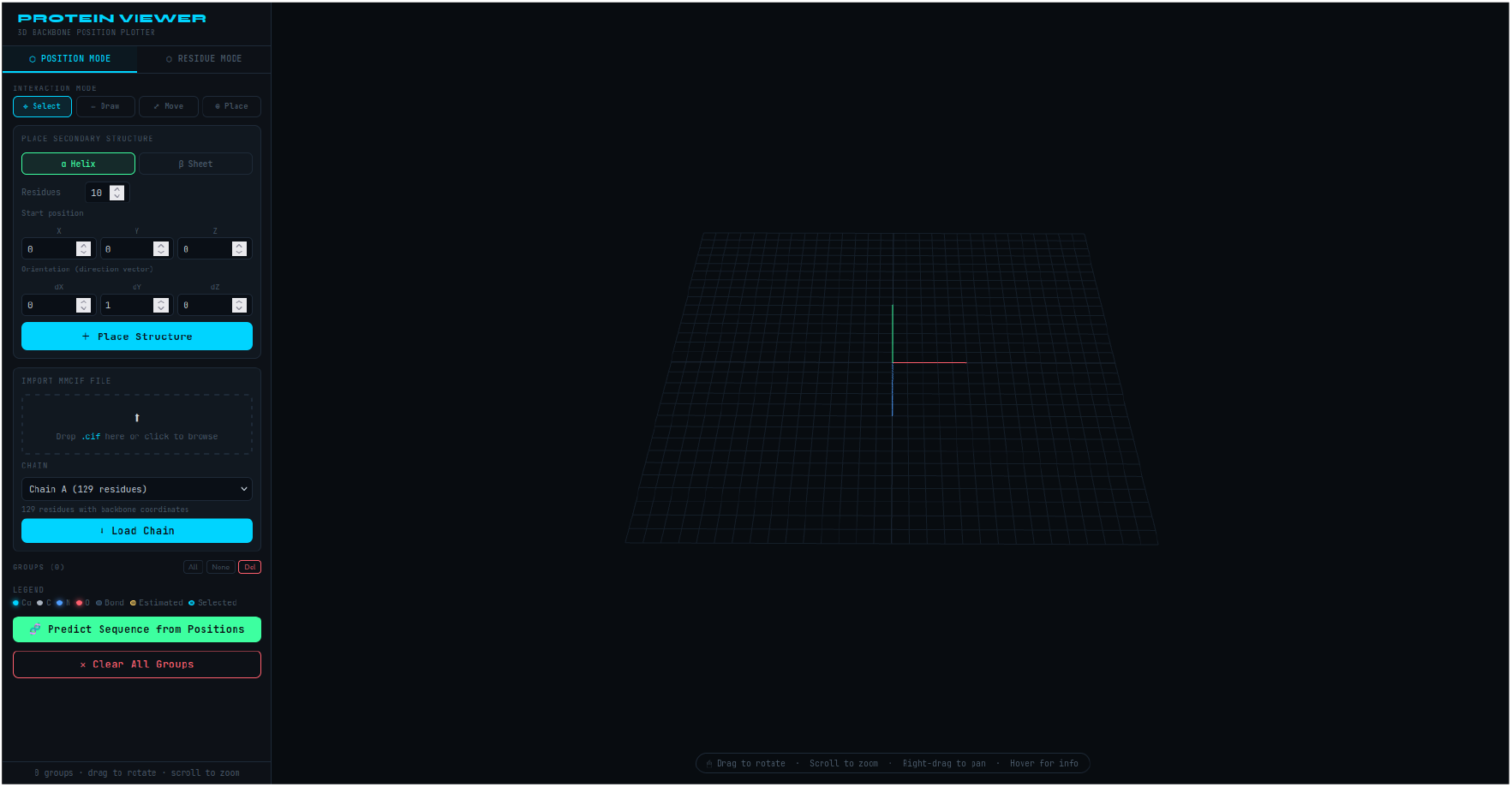
Overview of the interactive viewer. Left: the control sidebar, shown here in Position Mode. Right: the 3D viewport rendering a backbone with color coded backbone atoms (N, C*α*, C, O).

The rest of this paper surveys existing related tools (Section 2), describes the system’s architecture (Section 3), as well as its interface and functionality in the two modes, including a Kabsch alignment overlay tool for directly comparing two structures (Section 4), the workflow it enables (Section 5), a case study illustrating the workflow on an imported protein (Section 6), as well as its release as an open source code base which is intended for reuse (Section 7), finally closing with a discussion of its limitations and goals for future development (Section 8).

## 2. Related Tools

Some other general-purpose molecular visualization software, such as PyMOL [13] and ChimeraX [11], provide mature, high-accuracy renderings of protein structures. Still, neither is built around a sequence design model. Neither uses a point-and-click workflow to construct a hypothetical backbone from scratch or to immediately query a design model for a generated sequence and structure. Model-specific web servers, such as the public ProteinMPNN [2] server, execute one direction of the design problem for a fixed structure file, but do not provide backbone construction tools, nor do they close the loop by folding the resulting sequence back into a structure for visual comparison against the original design target. The tools in this paper are narrower in scope than general molecular viewers, but more interactive than fixed-model web servers; they also support hypothesis-driven experimental research and the models and proteins developed in our companion papers.

## 3. System Architecture

The system consists of two tools combined into a single interface: a local REST API that calls a Python/Flask backend hosting the trained models, and a single-file browser client (Section 3.2) that renders an interactive 3D scene with Three.js [14] and issues requests to the backend.

### 3.1. Backend

The Flask server loads two models at startup: ESMFold [8] and the structure-aware graph neural network described in our companion GNN paper [3]. This process requires three endpoints: predict sequence, predict structure, and server health.

- `POST /predict_sequence`: This endpoint takes in a list of backbone coordinates (N, C*α*, C, and O) and returns a predicted amino acid sequence, which was decoded randomly, together with per-residue confidence and the full 20 amino acid probability distribution at each position. This lets the user see confidence and other recommended amino acids.
- `POST /predict`: This endpoint receives an amino acid sequence and returns per-residue backbone coordinates and pLDDT confidence from ESMFold.
- `GET /health`: This endpoint reports server and device status and confirms the backend is reachable before submission.

Running the backend locally, rather than requiring a hosted service, makes the tool’s only hard dependency a GPU-equipped machine capable of loading ESMFold, while still allowing the server to exist if necessary. The backend is built on PyTorch [10], the HuggingFace transformers library [15], the fair-esm implementation of ESM [12], and Flask [9].

### 3.2. Frontend

This client is a single, self-contained HTML file with no required build step: it uses Three.js loaded from a CDN for 3D rendering and base JavaScript for the interaction logic. It also includes an in-browser mmCIF parser (described in Section 4.2) that requires no server call. This design choice lowers the barrier to running and editing the code; if the server is run, it can be used anywhere with the server URL.

## 4. Interface and Functionality

### 4.1. 3D Viewport

The viewport renders the protein backbone geometry with WebGL via Three.js. The atoms in amino acids are rendered as color-coded spheres (C*α* in cyan, N in blue, C in gray, O in red) connected by backbone binding lines. A trace connecting consecutive C*α* residues is overlaid on top of the amino acid structure to aid structural interpretation. The viewport supports standard rotational navigation: left-drag to rotate, scroll to zoom, and right-drag to pan. It also supports dynamic grid scaling and axis indicators, which adapt to the size of the loaded structure. Hovering over any atom displays a tooltip showing the atom type, residue identity, three-dimensional coordinates, and pLDDT confidence. Sharp-eyed biologists may have realized that the lines that connect the C*α* are a gross representation of the bonds between amino acids, but they do not have an impact on the actual rotational outputs of the models, and it was decided to make it easier to visualize.

### 4.2. Position Mode: Manual Backbone Construction

Position Mode lets users construct a backbone from scratch by placing residues (N, C*α*, C, O) into the 3D scene of Section 4.1. To make this easier, we provided the user with four interaction tools (when referring to these tools and structures, it is important that the reader and user understand that there is no placing of amino acids, only of the residues which are expected to be found in every amino acid group):

#### Place Tool

Residue groups can be placed by clicking in the 3D viewport, with the C*α* position anchored to a user-selected plane (XZ, XY, or YZ at a fixed depth). The C, N, and O atom positions are estimated automatically from the C*α* coordinate using a geometry-based estimation function. Afterward, the user can rotate the placed residues as they please. Alternatively, users can enter explicit Cartesian coordinates for all four backbone atoms manually; however, unless referencing an explicitly known structure, it is not recommended as the distances between residues are almost always the same.

#### Draw Tool

Clicking and dragging across the viewport creates a backbone continuously as the cursor is dragged along, placing new residue groups at a user-configurable minimum spacing (default 3.8Å, matching the standard C*α*–C*α* distance).

#### Secondary Structure Generator

Idealized alpha helices and beta strands can be placed directly at a desired location and orientation for a desired length. It connects to the most recently placed amino acid in the sequence. The alpha helix generator uses a rise of 1.5Å per residue, a radius of 2.3Å, and a rotation of 100° per residue. The beta strand generator uses a rise of 3.5Å per residue with alternating *±*1Å pleating perpendicular to the strand axis. Figure 2 shows both generators at their default parameters, an idealized helix in Figure 2a and a single strand in Figure 2b, each paired with the structure predicted from the sequence the GNN designs for it. Note that the beta strand generator places a single beta strand at a time: a beta sheet is not a stable structure on its own and only forms when multiple such strands are hydrogen bonded together side by side, so producing an actual sheet requires several strands positioned adjacently, most likely with the Select and Move tools described next.

**Figure 2.**
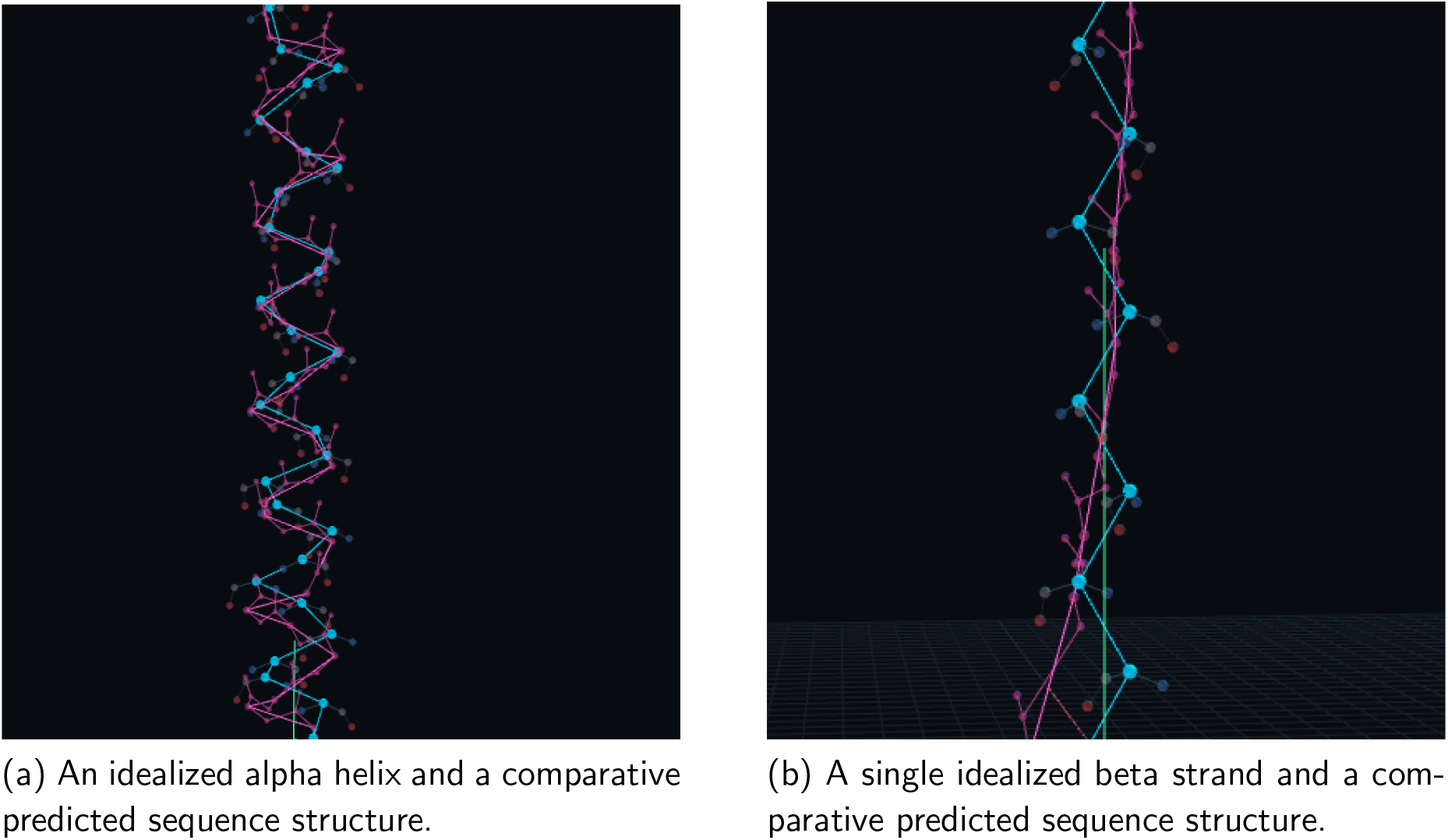
Backbones placed directly by the secondary structure generator, shown here with default parameters and no further editing. With the model that predicts a sequence and then compares the ideal structure to the model’s predicted structure. Panel (b) shows one strand in isolation; a real beta sheet is not a stable structure on its own and requires multiple strands hydrogen-bonded together, which the generator does not construct automatically. Hence the inability for the model to make a solid prediction.

**Figure 3.**
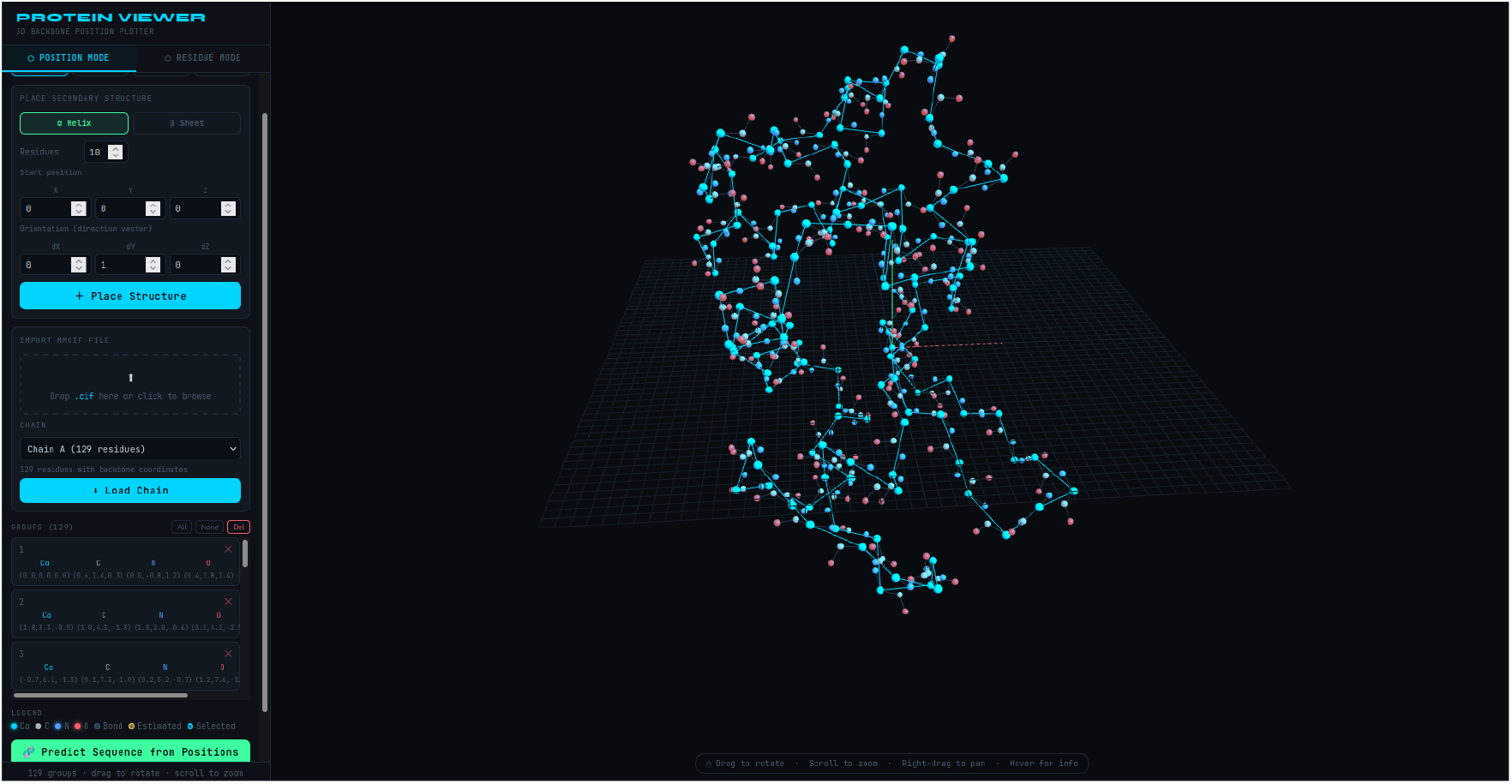
An example protein chain loaded into Position Mode via mmCIF import. This structure is used as the imported backbone example throughout Section 6.

#### Select and Move Tools

You can select individual residues or groups of residues and reposition them using point-and-click, a drag box, or a sequence of residues. Selected groups can be moved along the X, Y, or Z axes in configurable distances or rotated about any axis around their center, where their center is the average location of the C*α*.

#### mmCIF Import

Users can import experimentally determined, computationally predicted, or hand-designed structures directly from mmCIF files [1] by drag and drop or via the file browser. The parsing code scans and extracts backbone atoms across all chains, supporting multi-chain structures and allowing per-chain selection. This allows PDB structures to be loaded and used as backbone scaffolds for sequence prediction.

Once a backbone has been constructed or imported, clicking “Predict Sequence from Positions” submits the group coordinates to the /predict sequence endpoint. The GNN performs random decoding and returns the predicted amino acid sequence, which is then displayed with the per-residue confidence bars, as shown in Figure 4.

**Figure 4.**
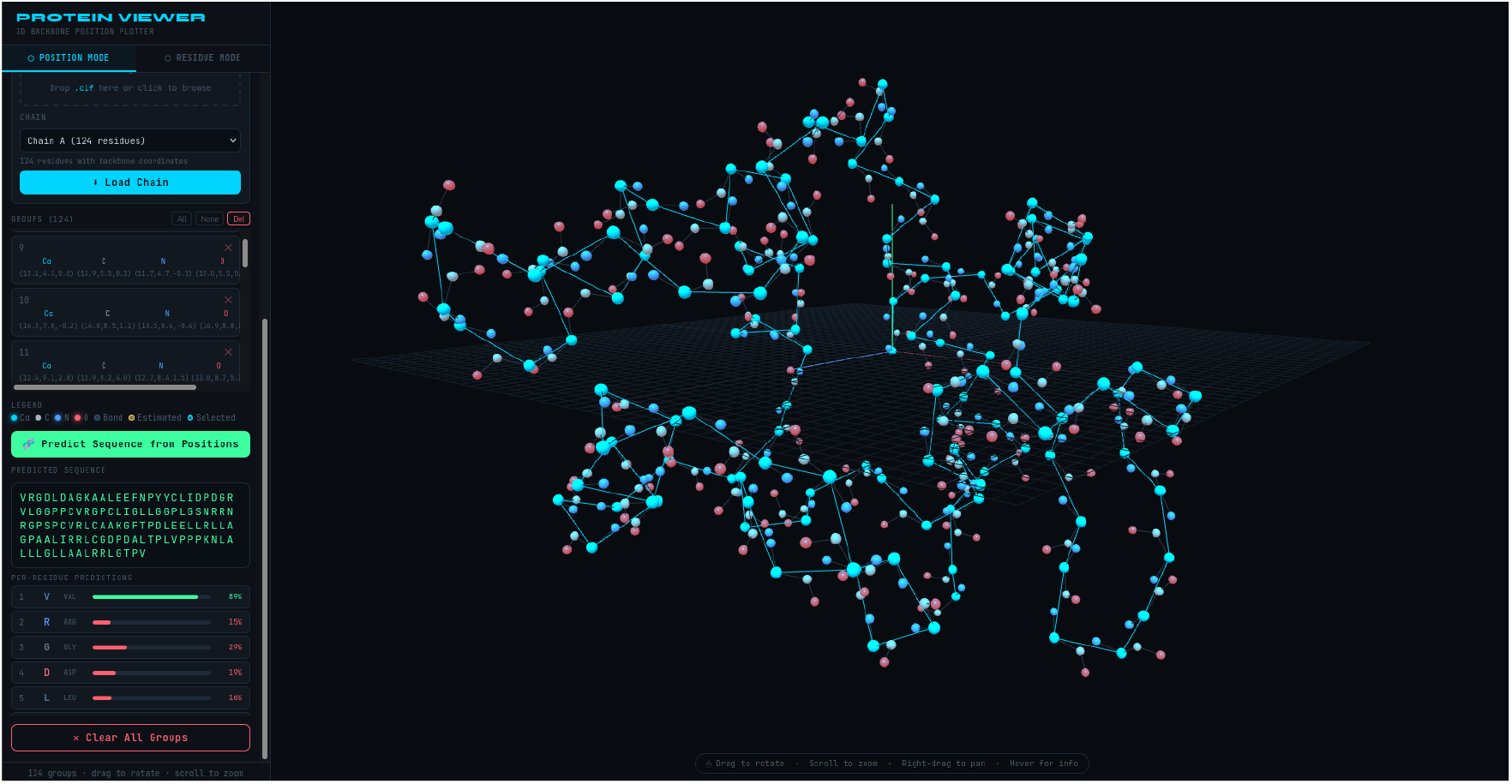
Predicted sequence panel following a /predict sequence call, displaying the decoded sequence and per-residue confidence bars on the left.

### 4.3. Residue Mode: Sequence to Structure Prediction

Residue Mode performs the inverse of Position Mode (Section 4.2): the user enters an amino acid sequence in single-letter code, easily taken from the sequence panel of that mode, using the /predict endpoint and folds it into a three-dimensional structure via ESMFold. The resulting backbone is then rendered in the viewport. Three atom coloring schemes are available: by residue, by per-residue pLDDT confidence (color-coded from high confidence[blue] to low confidence [red]), and by physicochemical amino acid groups (this is mostly meant to help with decoding of bad predictions). Figure 5 shows the pLDDT coloring applied to a predicted structure.

**Figure 5.**
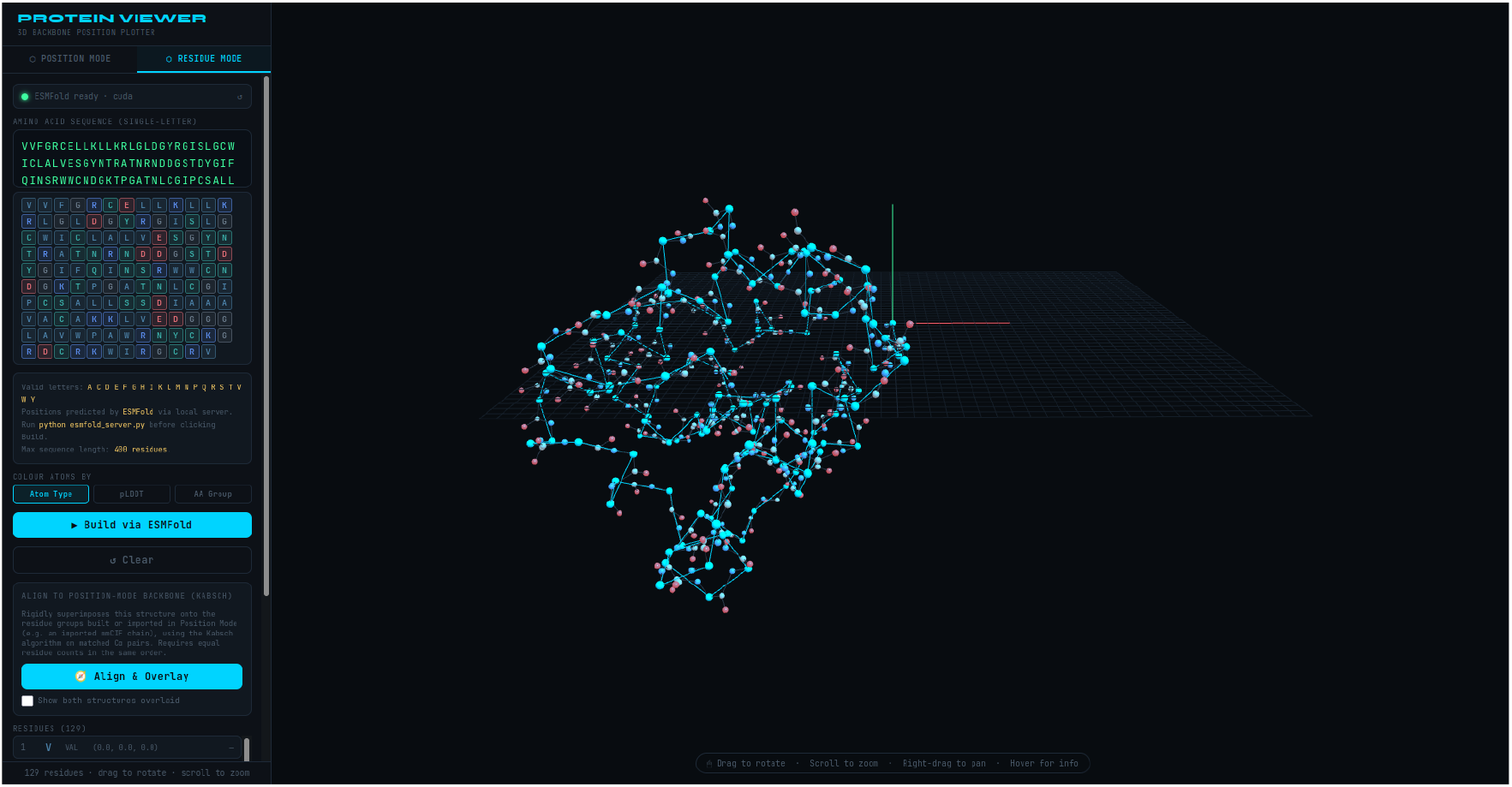
Residue Mode, showing an ESMFold-predicted structure colored by per-residue pLDDT confidence of the predicted sequence.

### 4.4. Kabsch Alignment and Structural Overlay

While a backbone built or imported in Position Mode and a structure folded in Residue Mode are by default translated independently so that their first residue C*α* sits at the origin, aligning the two structures is not very simple, and comparing them with the naked eye is unreliable, since what can be considered a bad residue prediction may only reflect orientation rather than structural differences.

To address this issue, the system implements Kabsch [6] superposition directly, requiring no server calls. Given the matched C*α* coordinates of the two structures, it computes the rigid-body rotation and translation that minimize their root mean square deviation, using the quaternion (Kearsley [7]) formulation of the Kabsch algorithm. A 4 *×* 4 symmetric matrix is constructed from the 3 *×* 3 cross-covariance matrix of the centered coordinate sets. Its dominant eigenvector, computed using cyclic Jacobi rotations, yields the optimal rotation in the form of a unit quaternion. This approach eliminates the need for a general-purpose SVD implementation in the browser.

Selecting “Align & Overlay” applies the resulting transformation to every backbone atom of the Residue Mode structure, then renders both structures together in the same view; the Position Mode backbone is displayed in its original coloring, and the aligned Residue Mode structure is shown in a distinct bright magenta, so the two remain visually distinguishable. The post-alignment C*α* RMSD is reported in the sidebar, providing quantitative data for the visual overlay. A checkbox can toggle the combined view on and off without recomputing the alignment so that the user can more easily spot the differences in structure; any action that changes one of the two structures will reset the alignment states. Additionally, the system requires that both structures contain the same number of residues.

## 5. Workflow

The two modes of Sections 4.2 and 4.3 support an end-to-end de novo protein design workflow:

1. The user constructs a target backbone geometry in Position Mode with any combination of manual placements, drawings, secondary structure generators, or mmCIF imports.
2. The backbone is sent to the GNN through the /predict sequence endpoint of Section 3.1, which predicts an amino acid sequence optimized for that geometry, using the input model.
3. The predicted sequence is then copied into the Residue Mode and pushed through ESM-Fold, which folds it into a predicted three-dimensional structure.
4. The Align & Overlay function aligns the ESMFold predicted structure onto the original backbone using the Kabsch overlay tool (Section 4.4) and compares the two structures, visually and quantitatively via the C*α* RMSD.
5. Based on the results, the user can iterate by modifying or re-predicting the sequence and repeating steps 3 and 4.

This closed-loop design, summarized in Figure 6, provides immediate visual and quantitative feedback on the structural quality of predicted sequences without requiring any external tools or command line operations.

**Figure 6.**
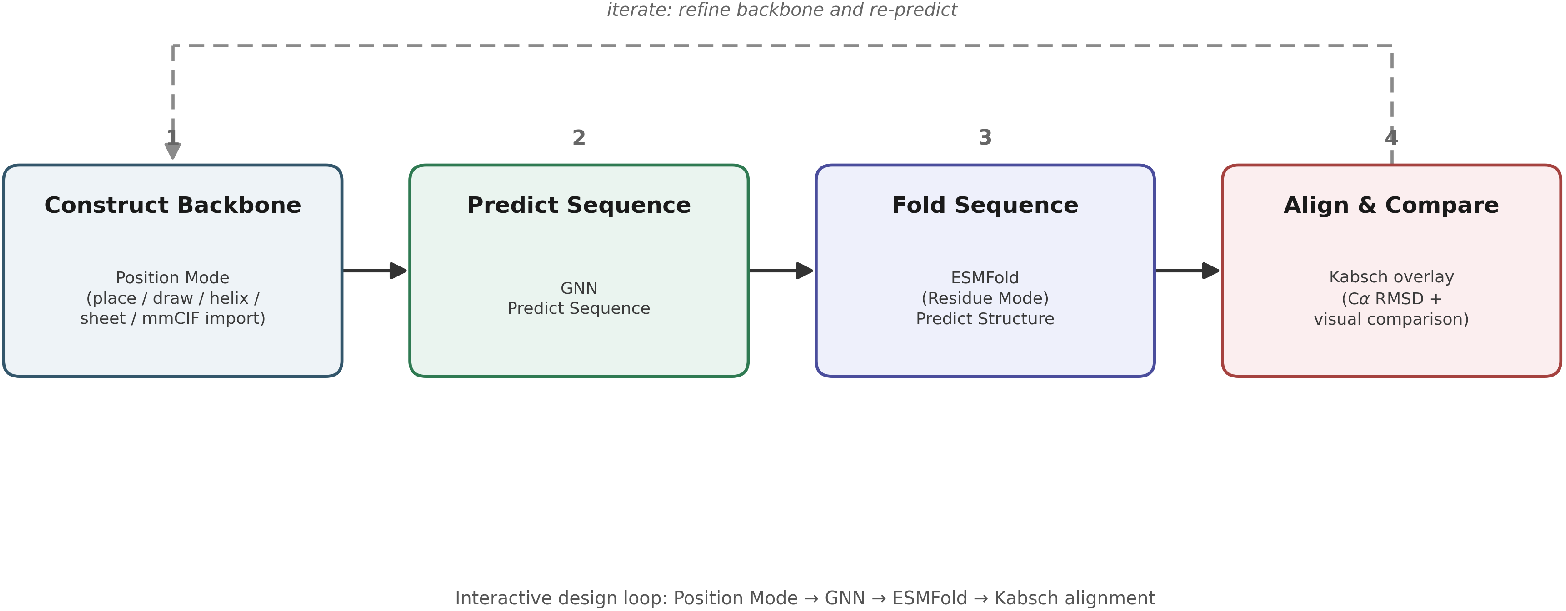
The closed design workflow for creating de novo proteins.

## 6. Illustrative Case Study

This section walks through an example of an imported real protein chain (Figure 3) and an example of evaluating the real sequence against the real structure. The purpose of this section is to show the interactive end-to-end workflow and to show that the tool surfaces enough information for a user to inspect and judge a single result themselves, which is the kind of use case this tool was designed for.

### 6.1. Round Trip from an imported structure

Starting from a backbone imported from an mmCIF file (Section 4.2, shown in Figure 7), the user can predict the sequence with the GNN model served by the backend of Section 3.1 (Figure 8). Copying the predicted sequence into Residue Mode and folding it with ESMFold yields the desired predicted three-dimensional structure of the predicted sequence. Aligning and overlaying the original structure with the predicted structure of the predicted sequence with the Kabsch tool from Section 4.4 shows how closely the ESMFold predicted structure recovers the original imported geometry (Figure 9). For this chain the round trip produced a post-alignment C*α* RMSD of 1.174Å.

**Figure 7.**
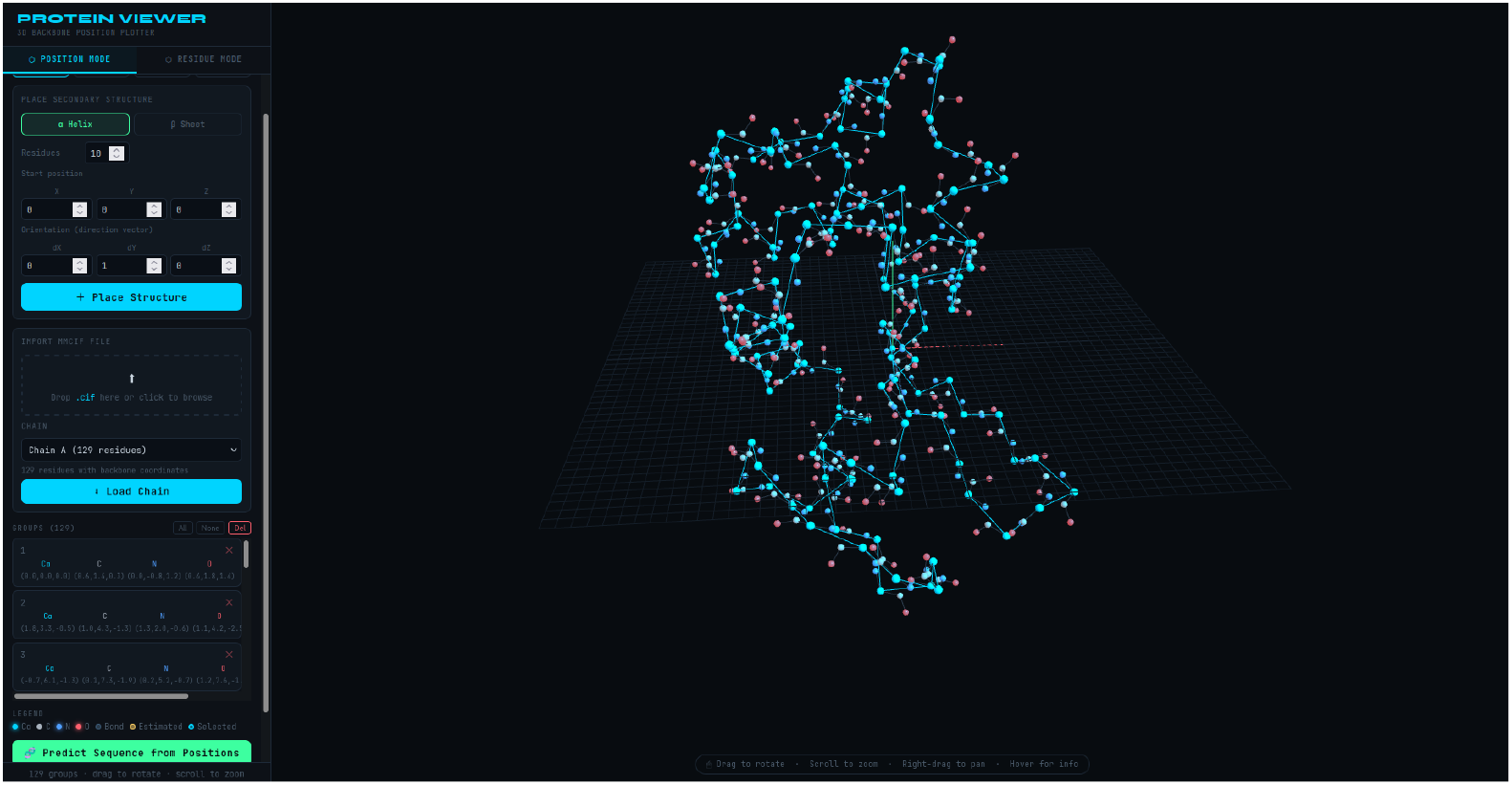
The imported mmCIF structure of the real protein.

**Figure 8.**
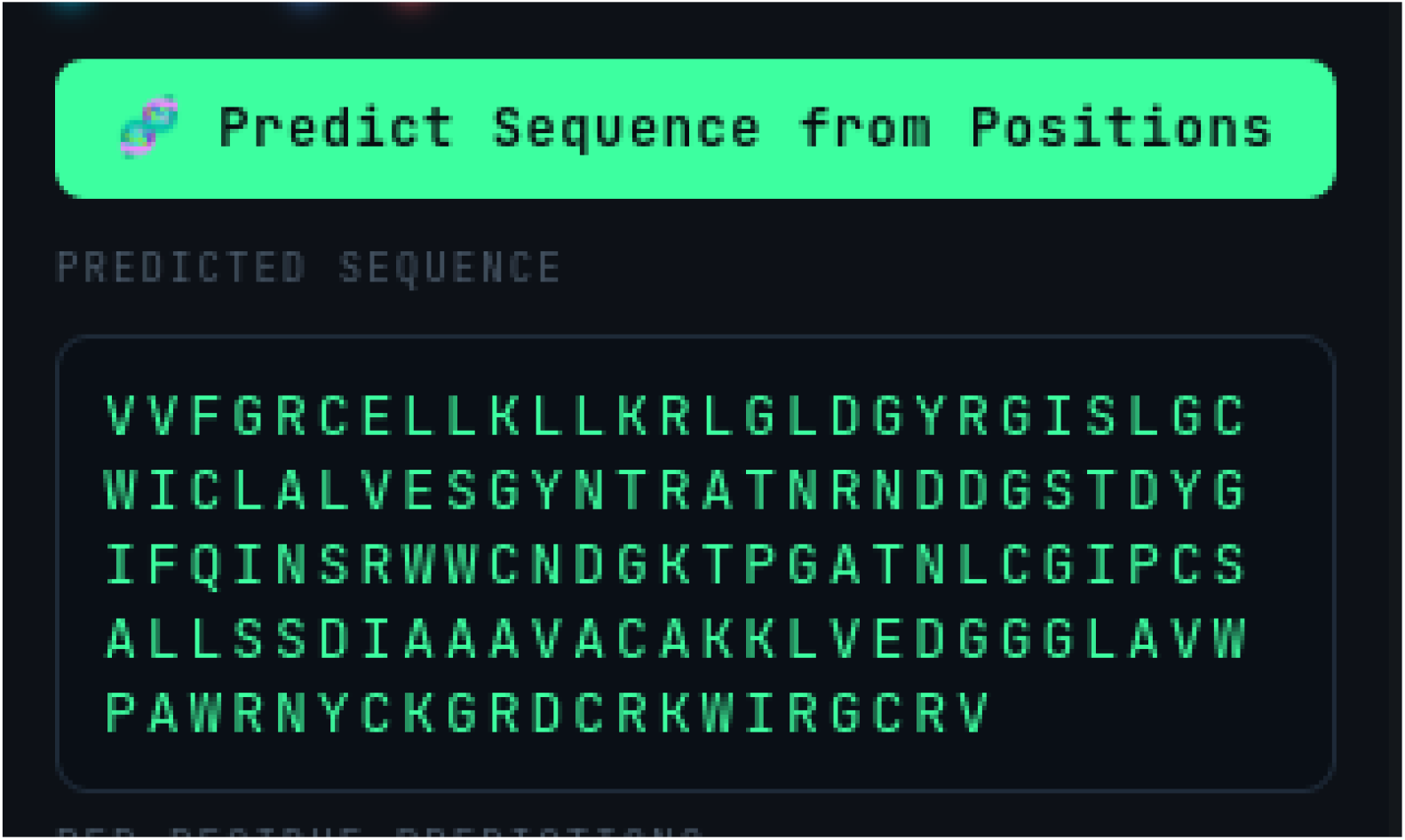
The GNN predicted sequence for the imported backbone of Figure 7.

**Figure 9.**
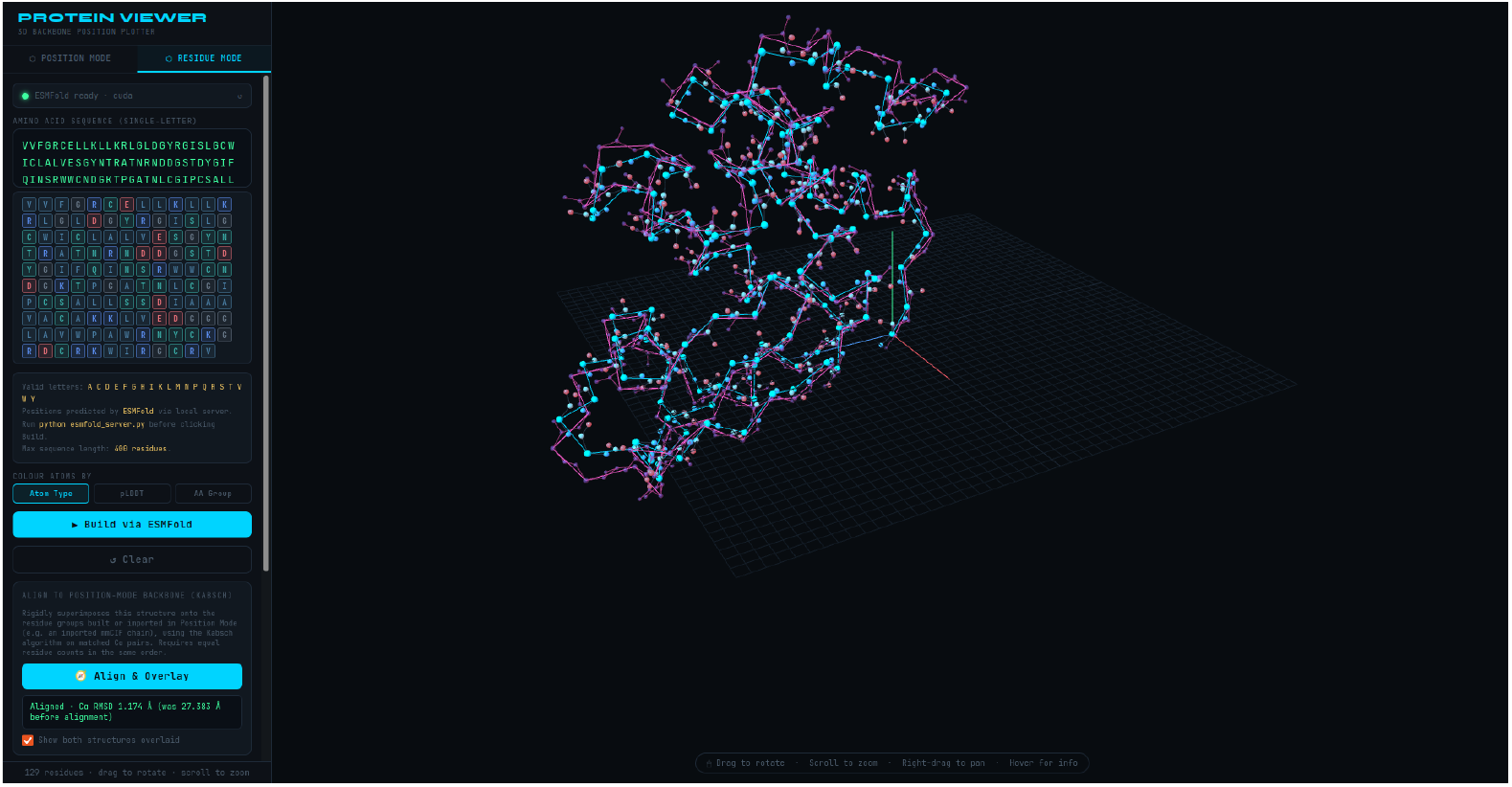
The overlay of the original imported structure and the ESMFold-predicted structure of the GNN-predicted sequence.

### 6.2. Validation: Refolding the Native Sequence

This second section tests the ability of the ESMFold model, which is a core model in this tool (Section 3.1), to properly reconstruct the “correct” sequence. To isolate how much residual error comes from the GNN’s sequence design versus from ESMFold’s own folding biases, the imported chain’s native sequence (not a GNN-predicted one) was entered directly into the Residue Mode, folded via ESMFold, and aligned back on top of the original imported structure. Figure 10 shows this overlay. Because both structures represent the same sequence, any remaining offset here reflects only ESMFold’s folding error, providing a baseline against which the GNN round trip overlay of Section 6.1 can be judged. For this chain that baseline was 0.999Å, against which the 1.174Å of the GNN round trip can be compared.

**Figure 10.**
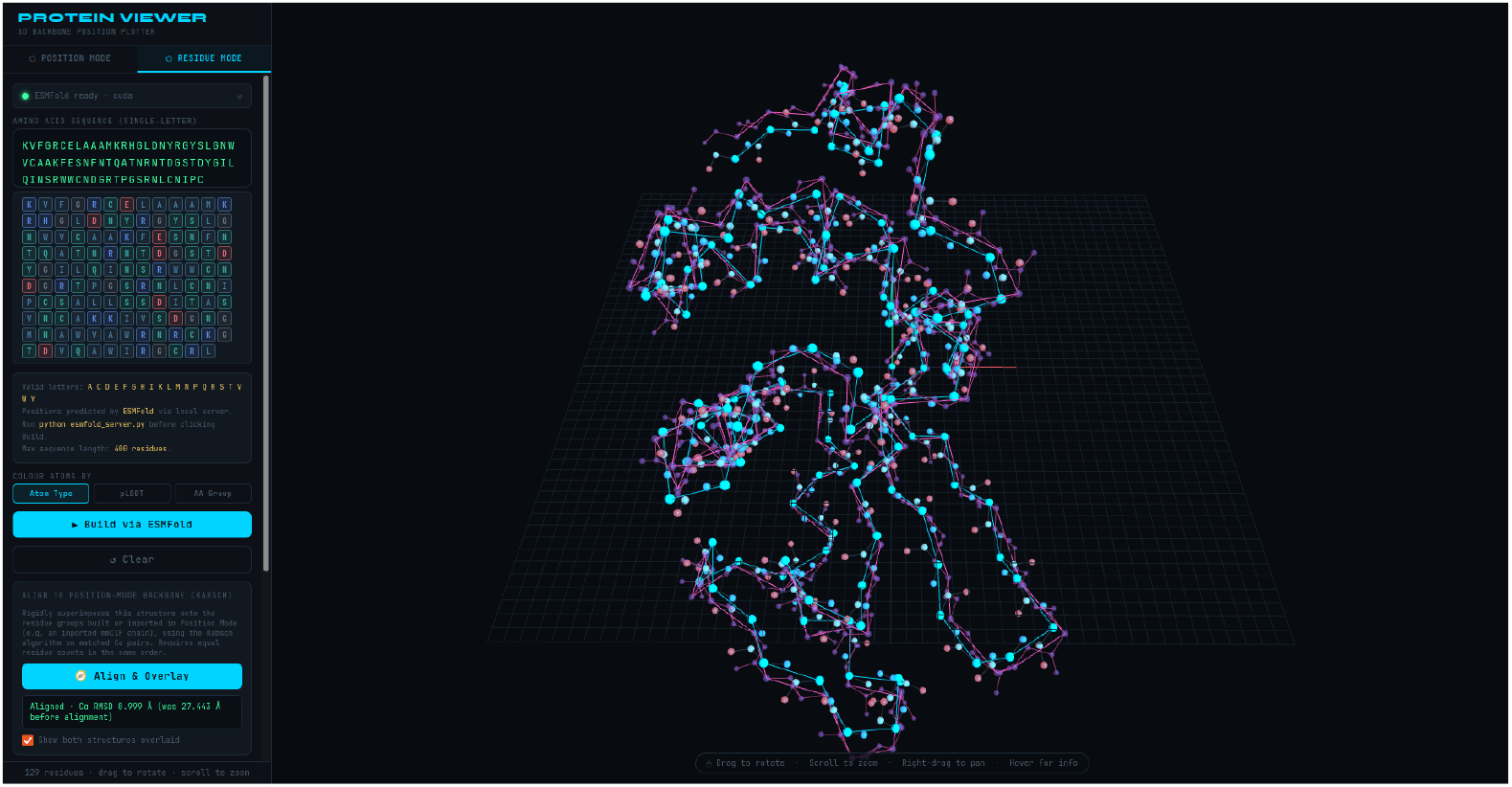
The imported backbone of Figure 7 aligned and overlaid against an ESMFold structure of the folded native sequence.

## 7. Open Source Availability and Reuse

The client and server code described in this paper are released under the MIT license and are intended for public release on GitHub, so that anyone can use, modify, and redistribute them freely. Two properties of the code implementation were prioritized specifically to support reuse by other researchers:

1. **Configurable backend** The GNN checkpoint path and architecture configuration are set by two variables at the top of the server script (CHECKPOINT PATH and MODEL CONFIG). Exchanging a different checkpoint trained with the same architecture, or including either of the two checkpoints produced for our companion papers [3,4], requires editing only CHECKPOINT PATH. Small changes to the models produced by our companion papers are possible by editing MODEL CONFIG; any changes where the model changes entirely must be performed by removing and importing one’s own model, though this requires more familiarity with our code base as well as the desired model’s code base. The intent is to let a group with its own trained checkpoints self-host an instance of the tool, rather than depend on a server maintained by the authors: running a persistent, GPU-backed public service carries an ongoing hosting and maintenance cost that this project does not currently plan to take on (see Section 8).
2. **Dependency-free client** As a result of the static single HTML front-end file, it can be adapted, embedded, or extended without a build toolchain, lowering the barrier for other researchers to modify the front-end interface for their own model or use cases.

## 8. Discussion

The protein design server demonstrates that the GNN model developed in our companion papers [3, 4] and ESMFold can be utilized in a practical interactive tool. By utilizing both models in a lightweight REST API and building a browser-based 3D interface, the system requires no installation on the user side, making it accessible to users without deep learning or programming expertise. The ability to import and quickly test mmCIF files is particularly useful for hypothesis testing, allowing a researcher to load a known protein structure, modify specific regions of the backbone, immediately query the GNN for sequences compatible with the modified geometry, and determine how likely it is to succeed, all in a closed workflow that would otherwise require multiple separate tools.

One of the largest limitations is that the server runs locally, which is a real barrier for users with little knowledge of programming. A persistent, web accessible deployment would remove that barrier, but was a deliberate non-goal of this tool, as the responsibility and cost of maintaining a server is disproportionate to the computational expenses of running the code. A low-cost, on-demand deployment, with a GPU instance that spins up on request rather than running continuously, would be worth exploring as a middle ground that decreases the hardware barrier without committing to a full-time server.

Additionally, the iterative refinement procedure described in our companion refinement paper [4] has not yet been integrated into the server interface, which, while significantly improving the quality of a prediction, was determined to be too time-consuming for the average prediction.

A further limitation is that the tool does not validate the input backbone geometry before submission. This means a hand-designed or edited structure with implausible bond lengths, angles, or steric clashes will still be passed onto the GNN, which returns a prediction regardless of how physically plausible the input was. This is a deliberate choice rather than an omission, allowing the user to ask for anything, and predicting as close as possible is the end goal of de novo protein generation. The interactive workflow is built around letting the user judge each result for themselves, so even if the per-residue confidence crashes, with confidence tending to drop in regions where the input geometry deviates from anything resembling a real protein structure, allowing the user to see a visual cue to revise the backbone rather than a validation error promotes de novo protein development.

The case study of Section 6 doubles as a sanity check on both models, comparing the GNN round trip of Section 6.1 against the native-sequence baseline of Section 6.2: the GNN-designed sequence and the refolded native sequence produced post-alignment RMSDs within 0.2Å of one another against the same imported backbone, indicating that both components operate as expected within the interface.

Finally, as one last check to determine the efficacy of this tool, a single 100-residue round trip was timed on the development machine (NVIDIA RTX 4080 Super GPU). The GNN’s /predict sequence call completed in 0.63s, and the ESMFold /predict call completed in 0.91s. As this was run locally, almost all of the wait the user experiences is model compute rather than network or serialization overhead. This is a single measurement at one sequence length on one machine; it is reported only to provide evidence of quick and useful predictions.

## 9. Conclusion

This paper discusses an interactive, open-source de novo protein design tool that unifies the structure prediction ESMFold model and the sequence prediction model developed in our companion papers [3, 4] into a single bidirectional design loop, accessible from a web browser with no software installation. Its value lies in closing the gap between the aggregate statistics used to evaluate sequence prediction models and what a researcher actually needs in order to judge a single design. A structure can score well on distance metrics such as RMSD, while still being blatantly wrong in some of the most important locations, and this tool is built to highlight those situations, both visually and quantitatively. As a preliminary check of the pipeline, a single case study achieved a GNN-predicted sequence RMSD of 1.174Å against the imported backbone structure, while the reconstruction via ESMFold of the “correct” sequence for the same protein’s native sequence achieved 0.999Å. This suggests that both ESMFold and the GNN model are performing adequately. Additionally, a single 100-residue round trip completed in under two seconds of compute, supporting the claim that the tool returns a usable estimate quickly enough for interactive use. In the companion refinement paper [4], it is mentioned that, at the cost of speed, the accuracy can be improved, but that was left as its own standalone code. The tools mentioned in this paper are released openly under the MIT license, are indifferent to which compatible checkpoint they serve, and are intended to let other researchers use the models described here interactively. It also encourages them to use the code as a front end for their own structure-conditioned protein sequence design work.

## 10. Declarations

### Funding

This work was supported by the Eugene McDermott Graduate Fellowship 202204.

### Use of generative AI

Claude (Anthropic) was used to assist with LaTeX formatting and editing. All content was reviewed, corrected, and finalized by the authors. The authors assume full responsibility for the generated content. No generative AI tool was used to produce, analyze, or interpret the experimental results.

### Competing interests

The authors declare no conflicts of interest.

### Author contributions

C.G.: conceptualization, methodology, software, validation, formal analysis, data curation, writing – original draft. L.T.: supervision, writing – review & editing.

### Data availability

PDB accession identifiers and evaluation splits will be deposited with the archived code release.

### Code availability

Model code, evaluation scripts, trained weights, and other code are available at https://github.com/Justworkname/GenomeGenerator

### Ethics approval

Not applicable; this study used publicly available protein structures and did not involve human participants or animals.

## Notes

https://github.com/Justworkname/GenomeGenerator

### Competing Interest Statement

The authors have declared no competing interest.

